# High δ^15^N values in Predynastic Egyptian archeological remains: A potential indicator for localised soil fertilisation practices in extreme conditions

**DOI:** 10.1101/2024.11.18.624066

**Authors:** Eve Poulallion, Annie Perraud, David Besson, Didier Berthet, Pascale Richardin, Arnauld Vinçon-Laugier, François Fourel, Jean-Pierre Flandrois, Bruno Maureille, Christophe Lécuyer

## Abstract

Predynastic Egypt arose around 5700 BC when nomadic people settled in the Nile Valley. Without voluntary practice related to anthropic mummification, as will be seen with the Egyptian Dynastic empires, human remains from this period were naturally mummified because of the natural environmental conditions, possibly enveloped in plant mats and animal skins. This study focuses on four mummies from the Natural History Museum of Musée des Confluences in Lyon (France). The geographic origin of the mummies and climatic conditions at that time were deduced from δ^18^O measurements on teeth and bones. These measurements confirm that the primary source of drinking water was the Nile, and that one mummy of unknown origin also originated from Upper Egypt like the other three. The diet and life habits of these individuals were inferred from carbon (δ^13^C), nitrogen (δ^15^N) and sulfur (δ^34^S) isotope compositions of their skins and bone collagens. Our study is of particular interest as we were able to analyse the animal skins and plant material enveloping the mummies using the same isotopic systems. The mean δ^15^N values obtained in human skins (17.9±2.4‰, AIR) were found to be high, and consistent with the values measured in animal skins (16.0±2.9‰, AIR). The analysed plants have even higher values, with an average of 22.1±2.2‰, AIR. The most probable explanation for the markedly elevated δ^15^N values observed in all the materials investigated is localised soil amendment practices. This is corroborated by the consistent nitrogen isotopic signatures across all the examined tissues, with a notable prevalence in plant tissues.

## Introduction

Understanding dietary practices is of great interest to advance global knowledge of the climatic and environmental conditions under which ancient civilisations flourished. Such practices also provide insight into human-induced modifications of ecosystems. Since the late 1970s, isotope geochemistry addressed these pivotal scientific questions in archaeology (Clauzel et al., 2020; Dodat et al., 2021; Jaouen et al., 2019; Vogel and van der Merwe, 1977; Yanes et al., 2011).

First, the study of food resources is a tool for the reconstruction of past climates with the measurement of oxygen isotopes (δ^18^O) in the phosphate group of hydroxyapatite in bones and enamel (Daux et al., 2008; Lécuyer et al., 1993). Ancient human populations consumed water from local springs, mostly replenished by rainwater, whose δ^18^O value depends on several isotopic latitudinal, altitudinal, continental, and seasonal gradients (Rozanski et al., 1993). In the specific case of Egypt, since the water initially comes from the Nile, the increase in δ^18^O in hydroxyapatite corresponds to the evolution of Nile δ^18^O, which records a global drying trend in the region (Touzeau et al., 2013). In addition, the carbon isotopic composition (δ^13^C) of collagen in bone remains provides insight into the types of plants consumed, namely C_3_, C_4_, or CAM, which thrive under different climatic conditions (Ambrose and DeNiro, 1989; Sternberg and DeNiro, 1983). This information provides an overview of the climatic conditions in the study areas over time. Consequently, the resources consumed, both food and water, provide insights into the climate, including both abundance and diversity of the food sources.

Secondly, the diet depends on socio-economic status of individuals at an archaeological site and the access to food resources including those derived from the marine environment. To study this, it is useful to examine the isotopic composition of nitrogen (δ^15^N), which provides insight into the diet of the subject, namely whether populations followed a vegetarian, omnivorous, or carnivorous diet with a specific ^15^N-enrichment in the case of marine food consumption (Herrscher et al., 2002; Katzenberg and Waters-Rist, 2018; Nehlich, 2015; Richards and Britton, 2020). In less wealthy populations, meat consumption is often limited or absent (Clauzel et al., 2023; Schwarcz and Schoeninger, 1991). To further specify the type of feed, the study of sulfur isotopic composition (δ^34^S) is useful for identifying potential marine resources (Bocherens et al., 2016; Britton et al., 2018; Ebert et al., 2021; Nehlich, 2015). In addition, the identification of isotopic signatures of non-local resources allows analysis of the commercial practices of individuals or groups, as demonstrated for Southeast Asia and the Middle East during the second millennium BC (Scott et al., 2021)

The study of the composition of the diets of past populations also involves the use of ecological engineering, essentially agricultural techniques that alter the ecology or composition of the soil and improve production yields (Schlütz et al., 2023). In Europe, livestock manure has been used for soil fertilisation and yield enhancement as well as water resource management since the Neolithic period (5900-2400 years BP) (Bogaard et al., 2013; Scharl et al., 2023). In South America, as early as the year 1000, seabird guano was used extensively as a fertiliser, particularly in the hyper-arid Atacama Desert (Santana-Sagredo et al., 2021). In Ancient Egypt, it is generally considered that the soil was, at least naturally fertilised when the Nile flooded, depositing fertile silt (El-Ramady et al., 2013; Hughes, 1992). Nevertheless, for some scientists (Y. Tristant, com. pers.), the annual deposit of Nile silts was no thick enough for fertilization and the need for organic natural amendment must have been significant.

The Predynastic Period in Egypt began around 5700 BC. It marked the transition from a nomadic to an agricultural society. This transition occurred when the population ceased migrations and settled along the banks of the Nile. They replaced their nomadic foraging practices with agricultural activities, including growing barley, flax, lentils, and onions, and raising cattle, ovicaprids, and goats (Lugan, 2021; Touzeau et al., 2014). This culture spread to several other sites, giving rise to several distinct predynastic cultures that lasted until about 3100 BC. The first settlements appeared during the Nagada II period (3500 to 3200 BC). During this period, collective works were established between the different populations on the banks of the Nile. Subsequently, the unification of Egypt took place gradually during the Nagada III period (3200 to 3100 BC), and the first dynasty emerged with the ascension to the throne of the first king of unified Egypt, Narmer (Lugan, 2021; Midant-Reynes, 2003, 1992; Shaw, 2003). During the Predynastic, it was common to bury individuals wrapped in cloth, animal skins, and plant mats (Crubézy et al., 2005, 2002; Holl, 2021; Janin, 1992; Midant-Reynes et al., 2002; Murray, 1956). These individuals were preserved by becoming naturally desiccated because of different environmental conditions (Jones et al., 2018).

Here, our focus is on the human remains of four Predynastic Egyptian mummies from the Musée des Confluences in France, home to a large Egyptology collection (Emmons et al., 2010; Madrigal et al., 2021). The four human mummies were buried according to the aforementioned methodology, without any additional embalming that was developed during the Old Kingdom period, which began around 2700 to 2500 B.C. (Abdel-Maksoud and Elamin, 2011; Jones et al., 2014; Nicholson and Shaw, 2000). As a result, these mummies are associated with elements of their environment, such as animal skins and plant fibers or textiles, and one of them was buried with polished stone tools models.

To determine the indigenous character of these mummies, the oxygen isotopic composition of hydroxyapatite found in human bones and teeth (δ^18^O_hap_) was first measured. The results of these analyses allowed us to determine whether a given individual had lived in an environment characterised by extreme heat and aridity, as occurred during the Dynastic Period (Touzeau et al., 2013). In order to study the diet of these predynastic Egyptians, the isotopic compositions of carbon (δ^13^C), nitrogen (δ^15^N), and sulfur (δ^34^S) were measured on their bone collagen and skin. The study of skin is particularly interesting because skin turnover is approximately 2% per day, documenting a period of up to a month and a half (Johns, 2012) and possibly two to four months (Lamb, 2016), allowing for the study of an individual’s diet over a relatively short period of time. This contrasts with the long-term averaging observed in bone collagen, which can span up to ten years, depending on the specific bone under consideration (Glimcher, 2006; Szulc et al., 2000). In addition to analysing the human remains, we performed the same isotopic measurements on the plant and animal skins enveloping the mummies. These data provided a more complete understanding of the environmental conditions and increased the reliability of our measurements on the human tissues. Considering the isotopic data obtained from all these tissues led to a new knowledge of the dietary habits of the Predynastic Egyptians and to consider the possibility of some specific ecosystem changes related to their agricultural practice in the context of a dry and hot environment.

## Material and methods

### 1. Material

The samples of mummies, animal skins and plants were obtained from the collections of the Natural History Museum of Musée des Confluences (France). These mummies are particularly rare, and our research team was fortunate to be granted permission to take samples from four of them. The rarity and interest of these samples is further enhanced by the fact that we have been able to sample the animal skins and plants that envelop the mummies. This allows, for the first time, a comprehensive multi-isotopic approach to be conducted on this set of samples. A photographic plate presenting the four mummies is shown in Figure 1.

**Figure 1:**
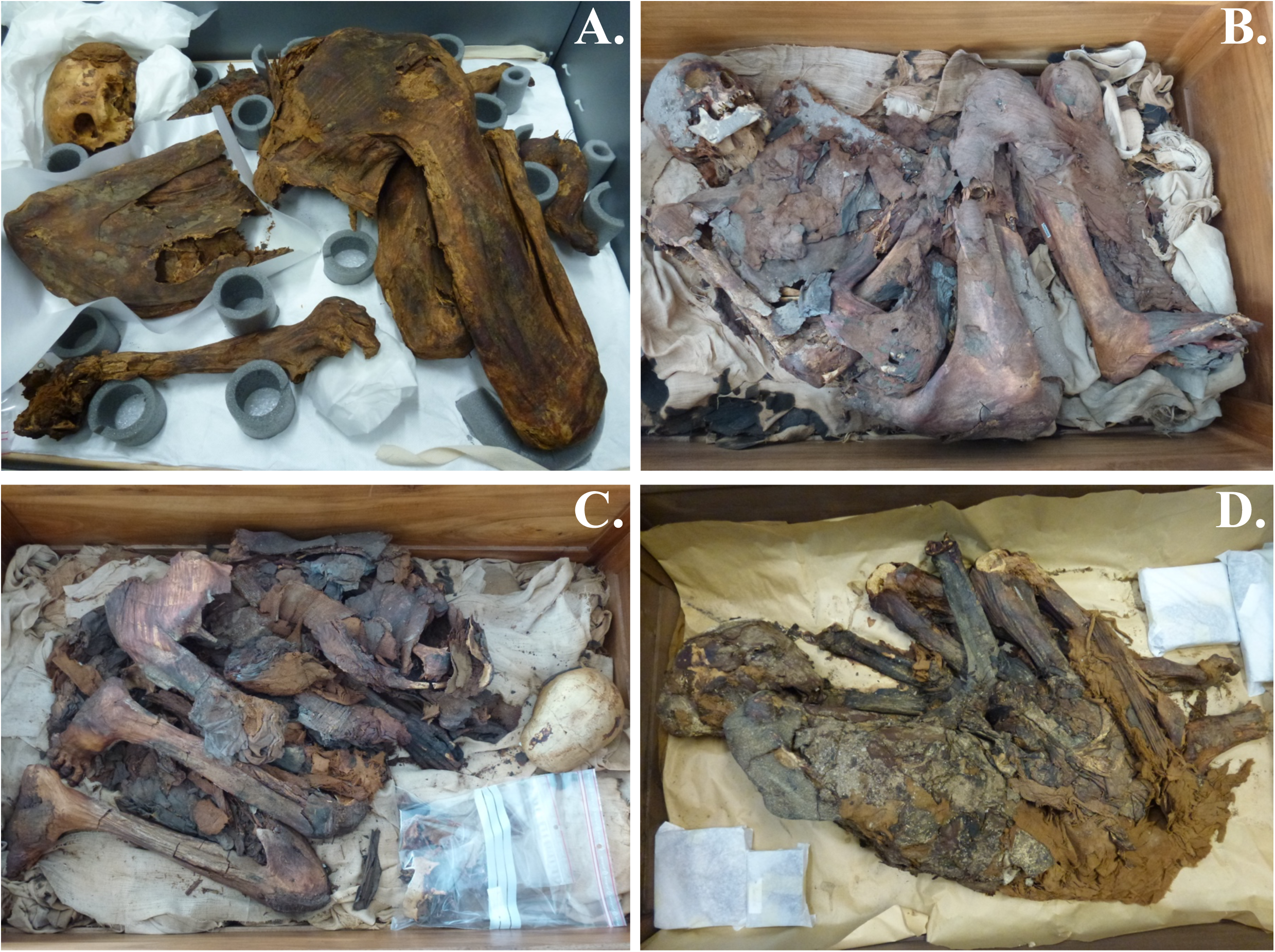
Photographic plate showing the four mummies from the Musée des Confluences: **A.** Mummy 90001635, **B.** Mummy 90002402, **C.** Mummy 90002403, and **D.** Mummy 90002404. Photographs were taken by A. Perraud.

Mummy 90002402 is that of an adult of undetermined sex, mummy 90002403 is that of a mature to elderly woman, and mummy 90002404 is a young adult woman. Finally, mummy 90001635 is that of a young adult woman. The mummies 90002402 and 90002403 are from Roda, while mummy 90002404 is from Gebelein. The provenance of mummy 90001635 remains uncertain. It should be noted, however, that all the Predynastic mummies in the museum originate from sites in Upper Egypt where excavation missions were conducted. A map is presented in Figure 2 to locate the different archaeological sites. Both identified archaeological sites are situated in proximity to the Nile Valley (Fig. 2). These two sites were ‘excavated’ by Charles-Louis Lortet (Lortet and Gaillard, 1909). X-rays of the lower extremities of the mummies 90002402, 90002404, and 90001635 revealed Harris lines on their skeletons. The presence of these lines on the skeletons of the mummies indicates that these human beings experienced a period of unspecific stress during childhood (Harris, 1933; Mays, 1995; McHenry and Schulz, 1976; Papageorgopoulou et al., 2011). Mummy 90002403 exhibits signs of meningeal infection, particularly periostitis on the bone of the skull. Individuals 90002402, 90002403, and 90002404 are from primary burials without skeletal remodeling. The type of burial for mummy 90001635, primary or secondary, is not entirely clear. The individuals were discovered in individual pits, covered with high-quality linen textiles, enveloped in a layer of animal skin and then of plant material. The mummy 90001635 was only covered by textiles and plant mat.

**Figure 2:**
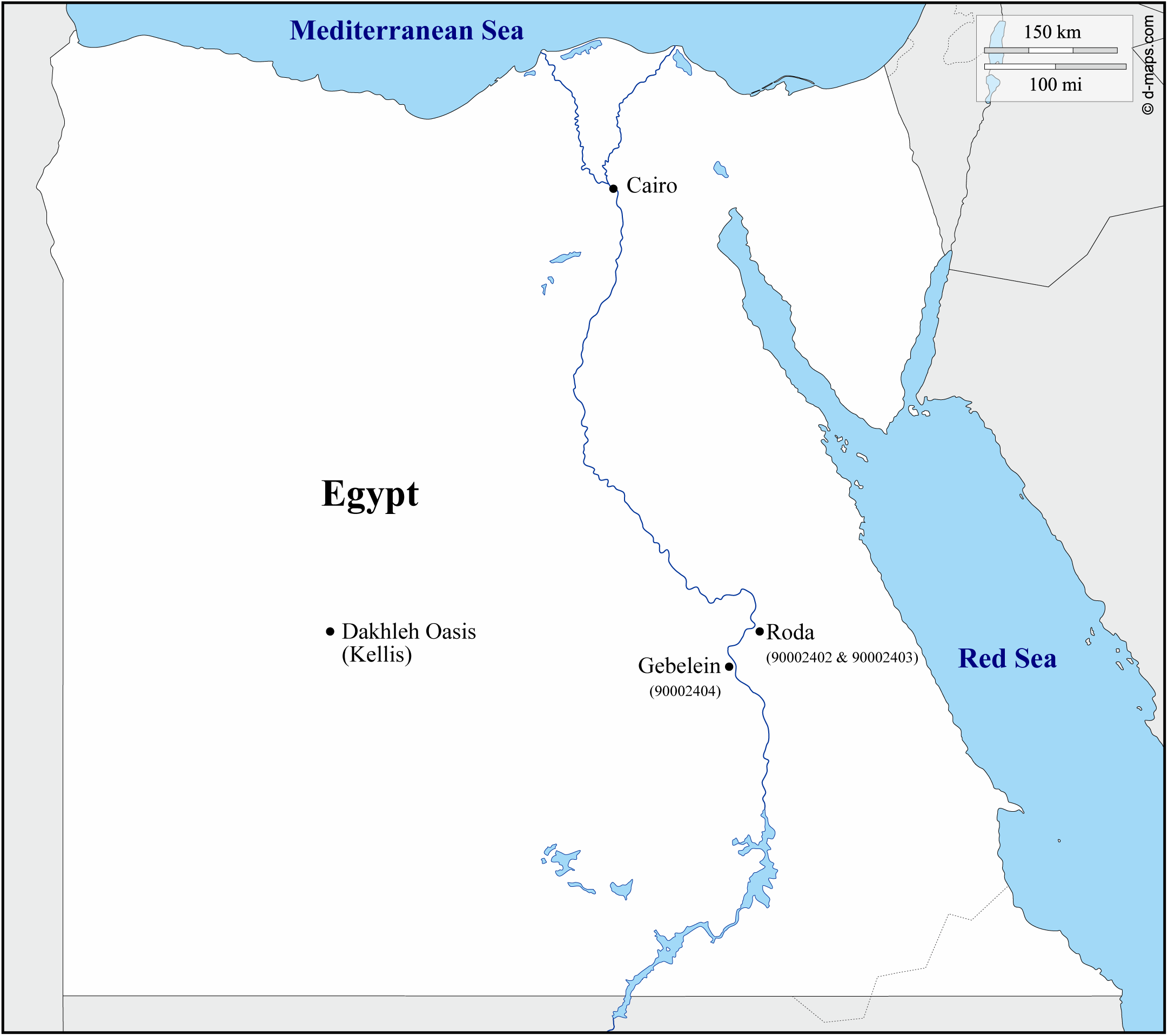
Map of Egypt, indicating the locations of the archaeological sites and the provenance of the mummies. The site of Dakhleh Oasis (Kellis) was also added to this map, as mentioned in the text as a mean of comparison. Map adapted from https://www.d-maps.com.

For the radiocarbon dating, samples of textiles were taken from each of the mummies, as well as a sample of plant material from mummy 90002404. For isotopic analyses, the skin samples were carefully removed with pliers. In order to avoid damaging the mummy, the selected bone samples were detached from the skeleton. For the 90001635 mummy, we collected a piece of the occipital bone and a first molar (M1). For the 90002402 mummy, we also collected an M1 and sampled the left ulna. For the 90002403 and 90002404 mummies, we collected pieces of the skull and the right tibia, respectively.

With regard to the animal skins, we proceeded to take one sample per human mummy, with the exception of mummy 90001635. The animal skins were meticulously separated from the mummies and removed with tweezers. The animal skins were identified as goat skins for mummies 90002403 and 90002404. The identifications were not possible for mummy 90002402. Hereafter, we will refer to these various skins as ‘animal skins’. With regard to the mats utilised to encase the mummies, we sampled plant material from the four distinct mummies. The plants used for the mats were identified at Xylodata in Paris (France) and were different for each mummies. *Desmostachya bipinnata* (L.) Stapf. Halfa. Poaceae was associated with every mummy, but mummy 90001635 also had *Juncus rigidus* Desf. Juncaceae, mummy 90002402 *Cyperus papyrus* L. Cyperaceae, and mummy 90002403 *Phoenix dactylifera* L. Arecaceae. We lacked sufficient material for the analysis of the plant material associated with the 90002404 mummy, which did not have another plant associated with *Desmostachya bipinnata* (L.) Stapf. Halfa. Poaceae

### 2. Methods

#### A. Radiocarbon dating

^14^C dating was conducted on tissue samples extracted from the mummies, resulting in a total of six samples. Additionally, a sample of the plant mat associated with mummy 90002404 was analysed. The process involved multiple steps, including two pretreatments. The first was a physical cleaning of the samples, and the second was a chemical extraction of carbonaceous material using the method by Richardin et al. (2017). The samples were then subjected to combustion. These procedures were conducted at the Centre de Recherche et de Restauration des Musées de France (C2RMF). The dating was performed using Accelerator Mass Spectrometry (AMS) in Saclay, France. Calendar ages were determined using OxCal v4.4 software (Bronk Ramsey, 2017, 1995; Bronk Ramsey and Lee, 2013) and the most recent calibration data IntCal20 (Reimer et al., 2020).The calibration results are considered within a two-sigma interval (i.e., a 95.4% confidence level) and expressed in terms of BC years.

#### B. Sample preparation

Mummies and animal skins were cleaned three times with a solution of dichloromethane and methanol (3:1 v/v) to remove mainly lipids. Following this process the skins were rinsed with deionised water, and then left to dry overnight in an oven at 45°C. The plant samples were cleaned five times solely with distilled water to avoid as the dichloromethane and methanol cleaning method might damage plant tissues.

The surface bone fragments of the four mummies were removed using a Dremel™ drill and an abrasive tip. The bones were rinsed with deionised water and subjected to ultrasonication to remove any residual internal residues such as soil. Following the cleaning process, the bones were dried in an oven overnight at 45°C and then ground into a fine powder.

#### C. Collagen extraction from human bones

The collagen extraction method was adapted from that described by Longin (1971) by Richardin et al. (2017). The bone powder was dissolved in 2 M hydrochloric acid (HCl) at a ratio of 500 mg to 5 ml. Subsequently, the solution was then diluted with 5 ml of deionised water and refrigerated for 5 hours. To neutralise the pH, the solution was washed several times and left overnight at room temperature. On the following day, 3 ml of 0.1 M sodium hydroxide (NaOH) was added, and the solution was allowed to react for one hour on ice. The solution was further washed to restore a neutral pH. An additional acidic treatment was conducted at 95°C by the addition of 4 mL of 2M HCl for 16 hours, resulting in the hydrolysis and dissolution of collagen within the solution. The solution was then filtered, freeze-dried, and the resulting dry collagen was stored in glass vials.

#### D. Silver phosphate precipitation from hydroxyapatite

The chemical procedure outlined by Crowson et al. (1991) and modified by Lécuyer et al., (1993) was employed to process human bones and tooth. Approximately 6 to 8 mg of powdered bone or tooth apatite were dissolved in 2 M hydrofluoric acid (HF) at ambient temperature for 24 hours. The resulting phosphate-bearing solution was then separated from the CaF_2_ precipitate via centrifugation. Subsequently, the CaF_2_ precipitate was triple rinsed with deionised water, with the resulting rinse water subsequently combined with the supernatant. The combined solution was then buffered with 2.2 mL of 2 M KOH solution. The neutralised solution was mixed with 2.5 mL of the cleaned Amberlite™ IRN78 anion-exchange resin. The mixture was agitated on a shaker table for a period of four hours in order to enhance the ion exchange process. The excess solution was discarded, and the resin was washed five times with deionised water to eliminate any residual contaminants. To quantitatively elute phosphate ions from the resin, the pH of the solution was diminished to 7.5 to 8.5 with 27.5 mL of 0.5 M NH_4_NO_3_ and the solution was agitated for four hours. The eluted solution was employed for the precipitation of silver phosphate crystals (Firsching, 1961). Afterwards, the solution was transferred to a 250-milliliter Erlenmeyer flask, where approximately 0.5 milliliter of concentrated ammonia hydroxide (NH_4_OH) was added to raise the pH to a range of 9 to 10. Then, a solution of ammoniacal silver nitrate (15 mL) was added to the flask and the solution was heated to 70°C in a thermostatic bath, resulting in the quantitative precipitation of millimeter-sized yellowish crystals of Ag_3_PO_4_. The Ag_3_PO_4_ crystals were collected on a Millipore filter, washed three times with deionised water, and oven-dried at 50°C.

#### E. Stable isotope measurements

The oxygen isotope data were obtained at the Genesis Platform of stable isotope measurements at the University of Lyon (LGL-TPE, UMR 5276) according to a high– temperature pyrolysis continuous flow technique (Fourel et al., 2011). Silver phosphates were weighed and placed in silver foil capsules, which were then pyrolysed at 1450°C using an Elementar VarioPYROcube™ elemental analyser. The latter is connected to an Elementar IsoPrime PrecisION™ mass spectrometer in continuous flow mode. NIST SRM 120c (δ^18^O_VSMOW_ = 21.7±0.2‰, Halas et al., 2011; Lécuyer et al., 1993) and NBS127 (δ^18^O_VSMOW_ = 9.3±0.4‰, (Halas and Szaran, 2001; Hut, 1987) were analysed alongside with the samples. In order to ascertain that the chemistry is operating as intended, silver phosphates were prepared from standard NIST SRM120c according to the same procedure as the samples.

Carbon and nitrogen isotope analyses (δ^13^C_VPDB_, δ^15^N_AIR_) of human skin samples were also conducted at the Genesis Platform. The sample introduction system is an Elementar Vario MICRO Cube™ system (Elementar GmbH Germany) based on purge and trap technology in combustion mode. It is connected in continuous flow with an GV IsoPrime60™ isotope ratio mass spectrometer (Elementar UK Ltd). The percentages of C and N are given by the Thermal Conductivity Detector (TCD). Isotopic data (δ^13^C_VPDB_, δ^15^N_AIR_) were calibrated with international standards, namely IAEA-600 (δ^13^C_VPDB_ = -27.77±0.04‰, δ^15^N_AIR_ = 1.0±0.2‰, Coplen et al., 2006), sucrose IAEA-CH-6 (δ^13^C_VPDB_ = -10.45±0.03‰, Coplen et al., 2006), and ammonium sulfate IAEA-N-2 (δ^15^N_AIR_ = 20.41±0.12‰, Böhlke and Coplen, 1995). These standards were used to correct the percentages obtained by the TCD.

To ensure reproducibility of our measurements, we conducted duplicate δ^13^C_VPDB_ and δ^15^N_AIR_ analyses on the human skins samples at the LEHNA (University of Lyon, UMR 5023). In this laboratory, we can measure during the same analysis δ^13^C_VPDB_, δ^15^N_AIR_ and δ^34^S_VCDT_. We also measured there the isotopic compositions of plants and animal skins as well. The isotopic analyses were performed using an Isoprime100™ isotope ratio mass spectrometer interfaced in a continuous flow to a VarioPYROCube™ system (Elementar GmbH Germany) based on purge and trap technology in combustion mode. Three sample aliquots were loaded into tin capsules to obtain δ^13^C_VPDB_, δ^15^N_AIR_ and δ^34^S_VCDT_ values and C, N, S percentages in the same analysis (Fourel et al., 2014). δ^13^C_VPDB_, δ^15^N_AIR_ and δ^34^S_VCDT_ values were calibrated against international standards, caffeine IAEA-600, sucrose IAEA-CH-6, ammonium sulfate IAEA-N-2, but also silver sulfide IAEA-S-1 (δ^34^S_VCDT_ = -0.30±0.03‰, Coplen and Krouse, 1998) and barium sulfate IAEA-SO-5 (δ^34^S_VCDT_ = 0.5±0.2‰, Halas and Szaran, 2001). C, N and S concentrations have been measured by using the TCD of the Vario PYRO Cube™ elemental analyser (Fourel et al., 2014). The elemental composition was calibrated using Sorghum Flour B2159 from Elemental Microanalysis (COA 257737), Protein B2155 from Elemental Microanalysis (COA 114859).

## Results

### 1. Radiocarbon dating

Table 1 presents a summary of radiocarbon age and the calculation of calendar ages (after calibration) for the samples. The results are also illustrated in Figure 3. The textile of mummy 90001635 was dated to 4795±30 BP (Before Present), which corresponds to a date between 3640 and 3525 BC. The combined radiocarbon age obtained for the textile of mummy 90002402 was 4676±22 years BP, which is consistent with a period between 3520 and 3370 BC. Likewise, the combined 14C age of the textile of mummy 90002403 was 4901±22 years BP, which is indicative of a period between 3710 and 3640 BC. At last, the combined ^14^C age of the textile of mummy 90002404 was 4763±22 years BP, which corresponds mostly to a period between 3635 and 3515 BC.

**Figure 3:**
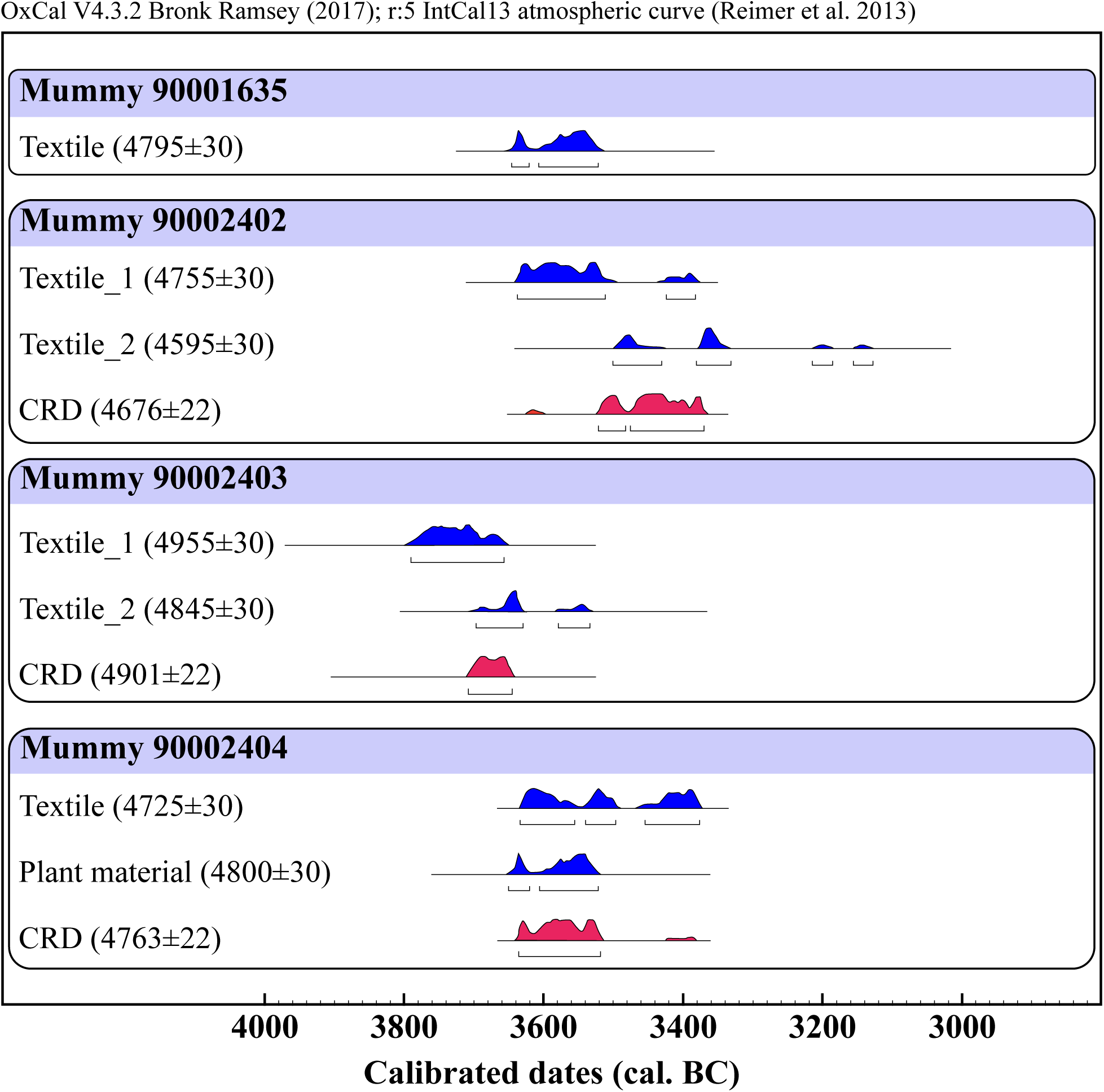
^14^C dating (Before Present) with corresponding calibrated dates at 95.4% probability of the Predynastic mummies of the Musée des Confluences. Data are presented under the form ‘sample type (^14^C date ± standard deviation)’ and calibrated dates are indicated at the bottom of the figure. Combined Radiocarbon Dates (CRD) are also given.

**Table 1:** Individual, sample type, and corresponding dates (at 95.4% certainty) inferred from ^14^C dating of the samples from the four Predynastic mummies from the Musée des Confluences. The Combined Radiocarbon Dates line (CRD) corresponds to the mean of the data measured and is checked for internal consistency by a chi-squared test which is performed automatically by the OxCal^TM^ program.

| Individual | Sample type | $^{14}\text{C}$ ages BP | Calibrated dates - $2\sigma$ (95.4%) |
| --- | --- | --- | --- |
| 90001635 | Textile | $4795 \pm 30$ | 3640 BC (95.4%) 3526 BC |
| 90002402 | Textile_1 | $4755 \pm 30$ | 3636 BC (84.6%) 3508 BC<br>3428 BC (10.8%) 3382 BC |
| | Textile_2 | $4595 \pm 30$ | 3504 BC (38.2%) 3430 BC<br>3380 BC (48.2%) 3330 BC<br>3216 BC (6.0%) 3186 BC<br>3153 BC (3.0%) 3122 BC |
| | CRD | $4676 \pm 22$ | 3520 BC (24.3%) 3483 BC<br>3476 BC (71.2%) 3372 BC |
| 90002403 | Textile_1 | $4955 \pm 30$ | 3790 BC (95.4%) 3658 BC |
| | Textile_2 | $4845 \pm 30$ | 3704 BC (5.3%) 3680 BC<br>3656 BC (52.7%) 3618 BC<br>3587 BC (37.4%) 3528 BC |
| | CRD | $4901 \pm 22$ | 3711 BC (95.4%) 3637 BC |
| 90002404 | Textile | $4725 \pm 30$ | 3630 BC (33.3%) 3556 BC<br>3538 BC (22.5%) 3494 BC<br>3458 BC (39.6%) 3376 BC |
| | Plants | $4800 \pm 30$ | 3650 BC (20.4%) 3621 BC<br>3606 BC (75.0%) 3522 BC |
| | CRD | $4763 \pm 22$ | 3636 BC (94.9%) 3516 BC<br>3391 BC (0.5%) 3386 BC |

### 2. Human samples

The collagen extractions yielded 0.45% for mummy 90001635, 2.05% for mummy 90002402, and 0.63% for mummy 90002404. When collagen was extracted from the bone of individual 90002403, no mass was recovered.

Table 2 presents the isotopic compositions of carbon, nitrogen, sulfur, and oxygen in bones and skins, as well as the molar ratios of C:N, N:S, and C:S in bone collagen. The molar ratios C:N of the bone collagen exhibit a range from 4.1 to 5.0. The mean N:S and C:S ratios were 47±24 and 217±120, respectively.

**Table 2:** δ^18^O_hap_, δ^15^N, δ^13^C, and δ^34^S values, as well as the weight proportions (wt%) of N, C, S of the bones, teeth and skins of the four human Predynastic mummies. The C:N, N:S and C:S ratios are given as a preservation indicator of the chemical composition of bone collagen. δ^18^O_dw_ is also displayed, calculated from the Daux et al. (2008) equation.

| Individual | Sample type | $\delta^{18}\text{O}_{\text{hap}}$<br>(‰, V-SMOW)<br>Mean ± SD | $\delta^{18}\text{O}_{\text{dw}}$<br>(‰, V-SMOW)<br>Mean ± SD | $\delta^{15}\text{N}$<br>(‰, AIR)<br>Mean ± SD | $\delta^{13}\text{C}$<br>(‰, V-PDB)<br>Mean ± SD | $\delta^{34}\text{S}$<br>(‰, V-CDT)<br>Mean ± SD | %N | %C | %S | C:N | N:S | C:S |
| --- | --- | --- | --- | --- | --- | --- | --- | --- | --- | --- | --- | --- |
| 90001635 | Bone | 19.7 ± 0.1 | -3.4 ± 1.3 | 13.7 ± 0.2 | -21.4 ± 0.0 | 10.8 ± 1.3 | 10.0 | 43.1 | 1.02 | 5.0 | 22 | 113 |
| | Tooth | 20.3 ± 0.1 | -2.5 ± 1.3 | | | | | | | | | |
| | Skin | | | 16.5 ± 0.1 | -21.4 ± 0.0 | 2.5 ± 0.1 | 14.0 | 45.3 | 0.6 | | | |
| 90002402 | Bone | 20.7 ± 0.1 | -1.9 ± 1.3 | 14.3 ± 0.1 | -21.7 ± 0.4 | 17.9 ± 1.7 | 12.40 | 43.3 | 0.6 | 4.1 | 46 | 189 |
| | Tooth | 20.6 ± 0.2 | -2.0 ± 1.3 | | | | | | | | | |
| | Skin | | | 17.3 ± 0.4 | -21.6 ± 0.2 | 2.9 ± 0.1 | 7.7 | 38.3 | 0.6 | | | |
| 90002403 | Bone | 20.4 ± 0.1 | -2.3 ± 1.3 | | | | | | | | | |
| | Skin | | | 21.3 ± 0.4 | -19.6 ± 0.1 | -9.8 ± 0.2 | 12.9 | 45.1 | 1.4 | | | |
| 90002404 | Bone | 20.6 ± 0.1 | -2.0 ± 1.3 | 12.1 ± 0.1 | -22.2 ± 0.6 | 25.2 ± 2.9 | 9.70 | 40.7 | 0.31 | 4.9 | 71 | 348 |
| | Skin | | | 16.3 ± 0.0 | -20.2 ± 0.2 | -1.2 ± 0.0 | 13.8 | 45.7 | 0.6 | | | |

As regards the δ^18^O_hap_ data obtained on the four individuals, one bone and one tooth were analysed for mummies 90001635 and 90002402, and a single bone was used in the case of mummies 90002403 and 90002404. For the 90001635 mummy, the occipital bone has a value of 19.7±0.1‰, and the first molar of 20.3±0.1‰. For the 90002402 mummy, the left ulna bone exhibits a value of 20.7±0.1‰, and the first left molar of 20.6±0.2‰. A value of 20.4±0.1‰ was measured on the bone of the individual 90002403, and 20.6±0.1‰ for the individual 90002404. Daux et al., equation (2008) was used to assess the δ^18^O of drinking water (δ^18^O_dw_) of these individuals. For the occipital bone of the 90001635 mummy, the δ^18^O_dw_ calculated value is -3.4±1.3‰ and for the M1 -2.5±1.3‰. The M1 of the individual 90002402 allowed us to calculate a value of -2.0±1.3‰, comparable to the value measured in the ulna of 1.9±1.3‰. The skull of the 90002403 mummy gave us a value of -2.3±1.3‰, and the tibia of the mummy 90002404 a value of -2.0±1.3‰.

The mean δ^15^N value for the three individuals from which collagen was recovered, namely 90001635, 90002402, and 90002404, is 13.4±1.1‰ (σ, n = 3), with a mean SD of 0.1. The δ^13^C values exhibited an average of -21.8±0.4‰ (σ, n = 3) and a mean SD of 0.4. The δ^34^S values range from 10.8 to 25.2‰, with a mean of 18.0±7.7‰ (σ, n = 3), and a SD that averages 2.0.

Finally, δ^15^N, δ^13^C and δ^34^S measurements were taken on the skin samples of all four mummies. The δ^15^N values range from 16.3 to 21.3‰ with a mean of 17.9±2.4‰ (σ, n = 4) and a mean SD of 0.3. For δ^13^C, the average value is -20.7±1.0‰ (σ, n = 4), ranging from -21.6 to -19.6‰. The mean SD is 0.1. With regard to δ^34^S values, they range from -9.8 to 2.9‰, and average -1.4±5.9‰, with an average SD of 0.1‰.

### 3. Animal and plant samples

The analysis of animal skins was conducted analogously to that employed for human skins. The results are displayed in Table 3. The δ^15^N values of animal skins range from 13.0 to 18.7‰ with a mean value 16.0±2.9‰ (σ, n = 3) and a mean SD of 0.2. For carbon, the δ^13^C values range from -22.5 to -14.3‰. The mean value of the measurements is -19.3‰ (σ, n = 3), with a mean SD of 0.2. The mean δ^34^S is 3.7±1.3‰ (σ, n = 4) and values range from 2.4 to 4.9‰. The SD averages 0.2 over these analyses.

**Table 3:** δ^15^N, δ^13^C, and δ^34^S values, as well as the weight proportions (wt%) of N, C, S of the plants and animal skins enveloping the human Predynastic mummies.

| Individual | Sample type | $\delta^{15}\text{N}$<br>(‰, AIR)<br>Mean ± SD | $\delta^{13}\text{C}$<br>(‰, V-PDB)<br>Mean ± SD | $\delta^{34}\text{S}$<br>(‰, V-CDT)<br>Mean ± SD | %N | %C | %S |
| --- | --- | --- | --- | --- | --- | --- | --- |
| 90001635 | Plant | 24.7 ± 0.4 | -25.0 ± 0.0 | 3.8 ± 0.4 | 2.4 | 45.5 | 0.2 |
| 90002402 | Plant | 20.8 ± 0.1 | -24.3 ± 0.1 | 3.5 ± 0.3 | 2.1 | 40.7 | 0.1 |
| | Animal skin | 16.3 ± 0.1 | -22.5 ± 0.1 | 3.8 ± 0.1 | 5.0 | 46.3 | 0.4 |
| 90002403 | Plant | 20.9 ± 0.1 | -21.5 ± 0.2 | 1.8 ± 0.6 | 2.4 | 42.0 | 0.1 |
| | Animal skin | 18.7 ± 0.2 | -21.1 ± 0.2 | 2.4 ± 0.3 | 4.6 | 43.3 | 0.3 |
| 90002404 | Animal skin | 13.0 ± 0.2 | -14.3 ± 0.3 | 4.9 ± 0.2 | 13.7 | 48.4 | 1.4 |

The results of plant measurements are presented in Table 3 as well. The δ^15^N values were measured between 20.8 and 24.7‰ with a mean at 22.1±2.2‰ (σ, n = 3) and a mean SD of 0.2‰. The δ^13^C values average -23.6±1.9‰ (σ, n = 3). They range from -25.0 to -21.1‰ and have an average SD of 0.1. Finally, the δ^34^S values range from 1.8 to 3.8‰. They average 3.0±1.1‰ (σ, n = 3) with an SD on average of 0.4.

## Discussion

### 1. Sample preservation and diagenesis

A chemical yield below 1% is generally considered to indicate poor preservation of collagen samples (Dobberstein et al., 2009). Moreover, the C:N molar ratios of our bone collagens are well outside the expected range 2.9 to 3.6 (DeNiro, 1985). Similarly, the N:S and C:S ratios of our collagen samples are also outside the expected ranges of 200±100 and 600±300 respectively (Nehlich and Richards, 2009). These molar ratios indicate that the bone collagen samples have been poorly preserved, which is consistent with the low collagen masses that were recovered after extraction. This is not unexpected, given that these individuals had not undergone any chemical treatment subsequent to their death. Therefore, the results obtained on bone collagen will not be discussed in the following sections. Inversely, hydroxyapatite is known to be a resistant mineral to diagenetic processes and provide reliable δ^18^O values despite various preservation states (Cox and Mays, 2000; Koch et al., 1992; Monnier and May, 2019). Although there are currently no established criteria for the preservation of skin tissue, desiccated mummies have been observed to have well-preserved dermal tissue (Johns, 2012; White and Schwarcz, 1994). Therefore, while remaining cautious about potential alterations to dermal tissue, it is possible to study the results obtained here, for both human and animal skins.

The plants enveloping the mummies are composed of an average of 43% of carbon and 2% of nitrogen which is the expected range for plants (Pate and Layzell, 1981). As with the skins, the plants studied here have undergone a similar drying process and this process was shown to have a negligible impact on their original isotopic composition (Ferrio et al., 2020; Metcalfe and Mead, 2019; Suh and Diefendorf, 2020; Szpak and Chiou, 2020). Consequently, we will consider those results, alongside the δ^18^O values of hydroxyapatite and skin samples, as reliable samples.

### 2. Oxygen isotopes

The δ^18^O_hap_ values of the bones and teeth of the mummies enabled us to trace the isotopic compositions of the water they drank (δ^18^O_dw_). For all the mummies, the overall δ^18^O_dw_ of the water consumed is similar, with and average value of -2.1±0.2 (n = 6). This is consistent with their discovery nearby archaeological sites in Upper Egypt. The isotopic values for the Nile at this time are close to the δ^18^O values measured from the mummies, estimated to be between -1 and -3‰ (Touzeau et al., 2013). It can be thus reasonably inferred that all these individuals consumed water from the Nile, and likely resided in the vicinity. They did not live in a generally very hot and arid climate, with increased evaporation of Nile water and increasing δ^18^O values of the latter (Lloyd, 1966).

### 3. Carbon isotope results

The ^14^C dating of the plant element enveloping the mummy 90002404 confirms that the plant mat was contemporary with the burial and therefore the mummification (Table 1, Fig. 3). δ^13^C values of plants confirmed the presence of C_3_ plants (Ambrose and DeNiro, 1986; Herrscher et al., 2002). However, two C_4_ plants were identified in the 90002402 mummy mat plant, whereas the measured δ^13^C value is representative of a C_3_ plant. It is therefore possible that other plants were present in the plant mat made for the burial, but may not have been present in the sample taken for plant identification. The δ^13^C measured on human skins all show the same trend with C_3_ plant consumption (Table 2). In general, the δ^13^C values measured in skin are similar or very slightly lower to those measured in bone collagen and can be interpreted in the same way (Finucane, 2007; Iacumin et al., 1996; Lamb, 2016). Conversely, skin collagen is enriched in ^15^N compared to bone collagen, but the magnitude of this enrichment is not constant (from 2 to 4‰ maximum on average, Finucane, 2007; Iacumin et al., 1996). Most studies on Ancient Egypt demonstrate that the Egyptians consumed a majority of C_3_ plants (e.g. Thompson et al., 2005; Touzeau et al., 2014). This result is consistent with what we observe. We note that the animal skin enveloping the individual 90002404 exhibits representative values for C_4_ plants that flourish in more arid environments (Table 3, Ambrose and DeNiro, 1986; Herrscher et al., 2002).

### 4. Food resources influencing δ^15^N values?

While the δ^13^C data provide consistent values with the expected diet behaviour of Predynastic populations living along the Nile valley, the δ^15^N values of human skins are high. Such values are usually associated with marine diets (Nehlich, 2015). However, here, we do not compile with such hypothesis as the δ^34^S values measured in human skin samples do not indicate such marine origin of the product as the values measured are too low (e.g. average marine modern fish δ^34^S value of 16.8±0.7‰, Nehlich, 2015, δ^34^S values of macroalgae around 15-22‰, Kejžar et al., 2021; Velasquez-Vacca et al., 2024). Our second hypothesis would be to include freshwater supply such as Nile perch in the food of the individuals. Nile perch are expected to have δ^34^S values between 7 and 12‰ (Touzeau et al., 2014, Fig. 4). This is the range we would expect from tissue measurements if these individuals consumed a lot of freshwater resources, as there is little isotopic fractionation between prey and consumer for sulfur isotopes (Nehlich, 2015). However, our δ^34^S values are much lower, and the δ^15^N values observed in these same Nile perch are, at most, approximately 11‰, a value that is lower than what we measured, even considering trophic enrichment (Touzeau et al., 2014, Fig. 4). Therefore, we can also withdraw the assumption of important freshwater fish consumption.

**Figure 4:**
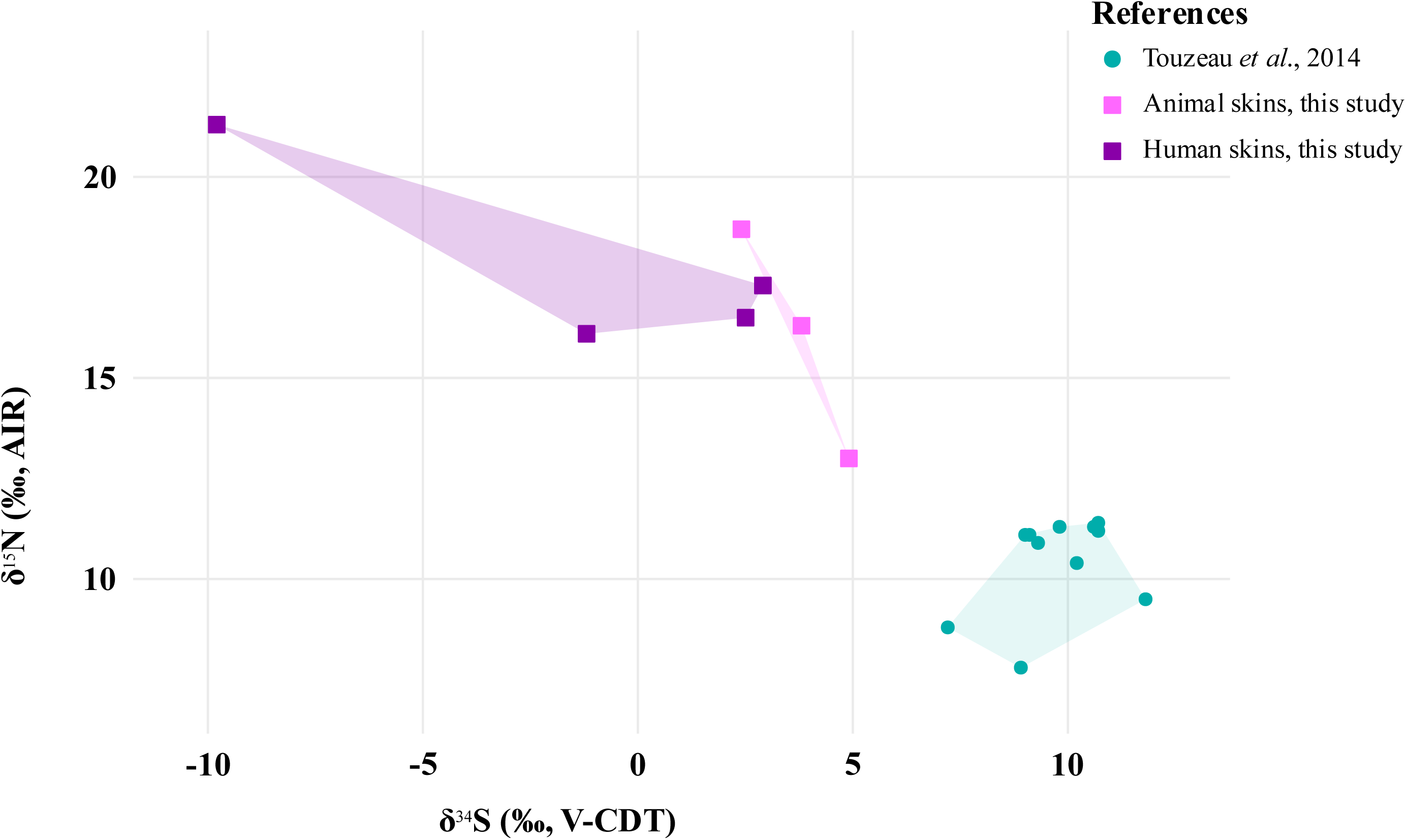
Nitrogen isotope compositions (δ^15^N) as a function of sulfur isotope compositions (δ^34^S) for animal and human skins of our study. Data is compared with δ^15^N and δ^34^S values published by Touzeau et al., 2014 for Nile perch of ancient Egypt.

### 5. δ^15^N values increased by aridity?

Such high δ^15^N values have previously been measured both in skins and bone collagens in Egyptian mummy samples, either associated with hyper arid environment or have been found with diseases (Fig. 5, Dupras and Schwarcz, 2001; Iacumin et al., 1998, 1996; Thompson et al., 2008, 2005). Soil ammonia is depleted in ^15^N compared to other organic nitrogen species. Ammonia volatilises from the soil as aridity increases (Schwarcz et al., 1999), impacting the entire food chain. For example, mummies from the Dakhleh Oasis, in the middle of the Egyptian desert (Fig. 2) possess high δ^15^N values of bone collagens (mean of 17.9±1.1‰, Dupras and Schwarcz, 2001, Fig. 5), and some of the individuals were supposed to have leprosis. None of the mummies from our study compile with one or the other of those criteria. Our δ^15^N values are also quite close to those measured in the skin of Egyptian and Nubian individuals published in previous studies (Iacumin et al., 1998, 1996), or even slightly higher (Fig 5). If we consider the mean offset of 3‰ between δ^15^N values of bone and skin collagens, we could conclude that the mean δ^15^N expected value for the bone collagen would be approximately 15‰. This mean value is close but slightly higher than the measurements made in previous studies on bone collagen (Iacumin et al., 1998, 1996; Thompson et al., 2008, 2005, Fig. 5). Thus, we propose that the human skins of our study are likely to be in a comparable state of preservation to those previously studied (Iacumin et al., 1998, 1996), given that they exhibit comparable δ^15^N values. The values published by Johns (2012, Fig. 5) are measurements on skin samples collected from Egyptian mummies of the Dakhleh Oasis, which was referenced above, and is distinguished by its extreme aridity. It is therefore unsurprising that the values measured by the latter are higher than ours, given that they reflect a different environment. In conclusion, the δ^15^N values measured on the individuals in our study are slightly higher than those of previously published studies, but lower than those typical of an extremely arid environment.

**Figure 5:**
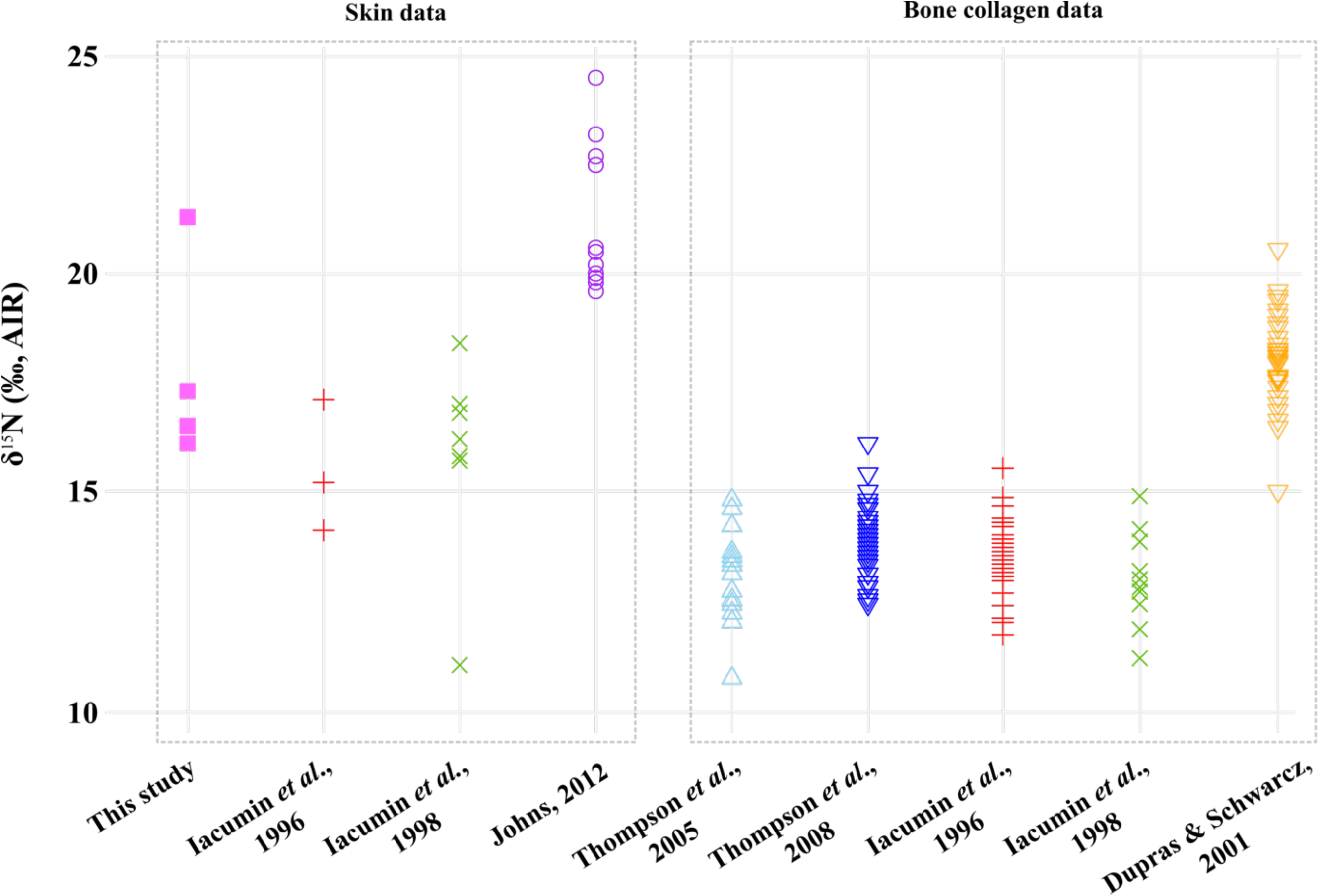
Nitrogen isotope compositions (δ^15^N) of human skins of our study compared with literature data of ancient Egyptians and Nubians (Iacumin et al., 1998, 1996; Johns, 2012). The data is also compared with δ^15^N values of bone collagens of Egyptians and Nubians measured in previous studies (Dupras and Schwarcz, 2001; Iacumin et al., 1998, 1996; Thompson et al., 2008, 2005).

We observe a lack of a trophic jump of +3.8±1.1‰ for δ^15^N values between animal skins and human skins (Bocherens et al., 2015), with the exception of mummy 90002403 (Table 2, Table 3, Fig. 5, Fig. 6). Animal and human skins δ^15^N values are similar, indicating that meat intake in the Egyptian diet was likely minimal, in accordance with previously published studies (Touzeau et al., 2014). The mean δ^15^N of animal skins is 16‰, or approximately 13‰ when the skin-collagen of 3‰ offset is applied. Our δ^15^N values are considerably higher than those previously published in the literature, considering a crocodile skin, a carnivorous animal, that was measured at 12‰ (Iacumin et al., 1996, Fig. 6) or bone collagens from other ancient Egyptian goats, which averaged 8‰ (Fig. 6).

**Figure 6:**
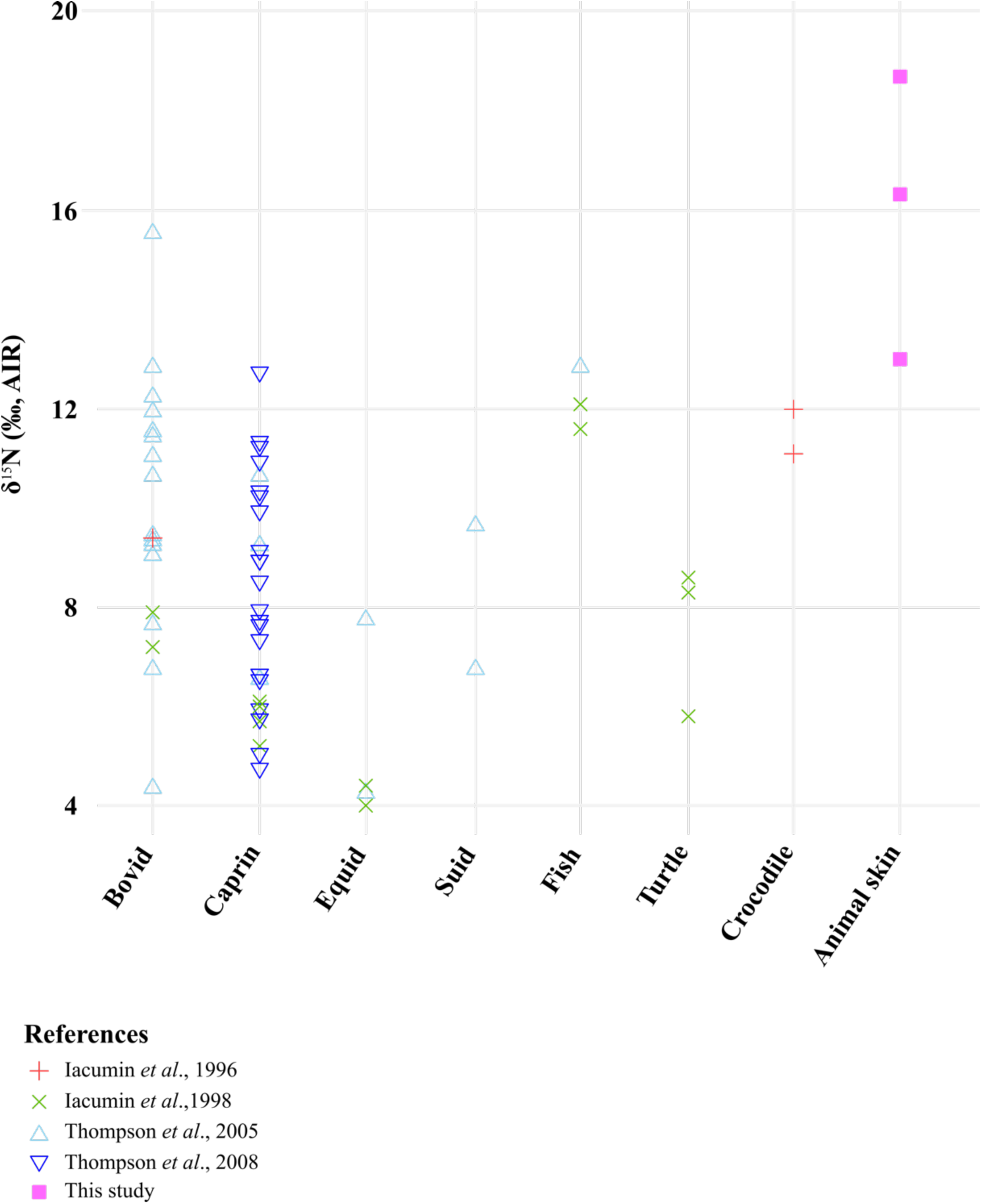
Nitrogen isotope compositions (δ^15^N) of animal skins of our study compared with previous published data for both skins and bone collagens of ancient and modern Egyptian and Nubian animals (Iacumin et al., 1998, 1996; Thompson et al., 2008, 2005).

Plants, which are at the bottom of the food chain, and reflecting the global environment, have δ^15^N values even higher values (approximately 22 ‰) than those observed in human and animal skin samples. Once again, our plant data are significantly ^15^N-enriched relative to previously published findings (Iacumin et al., 1998, 1996; Schwarcz et al., 1999, Fig. 7). Such isotopic pattern could be explained by a punctual episode of environmental aridity. Nevertheless, this phenomenon appears to be insufficient to account for the δ^15^N values that were observed. To fix ideas, an archaeobotanical study from the village of Kellis in ancient Upper Egypt (Dakhleh Oasis, Fig. 2) reveals δ^15^N values of plants to fluctuate between 12 and 19‰ (Schwarcz et al., 1999, Fig. 7).

**Figure 7:**
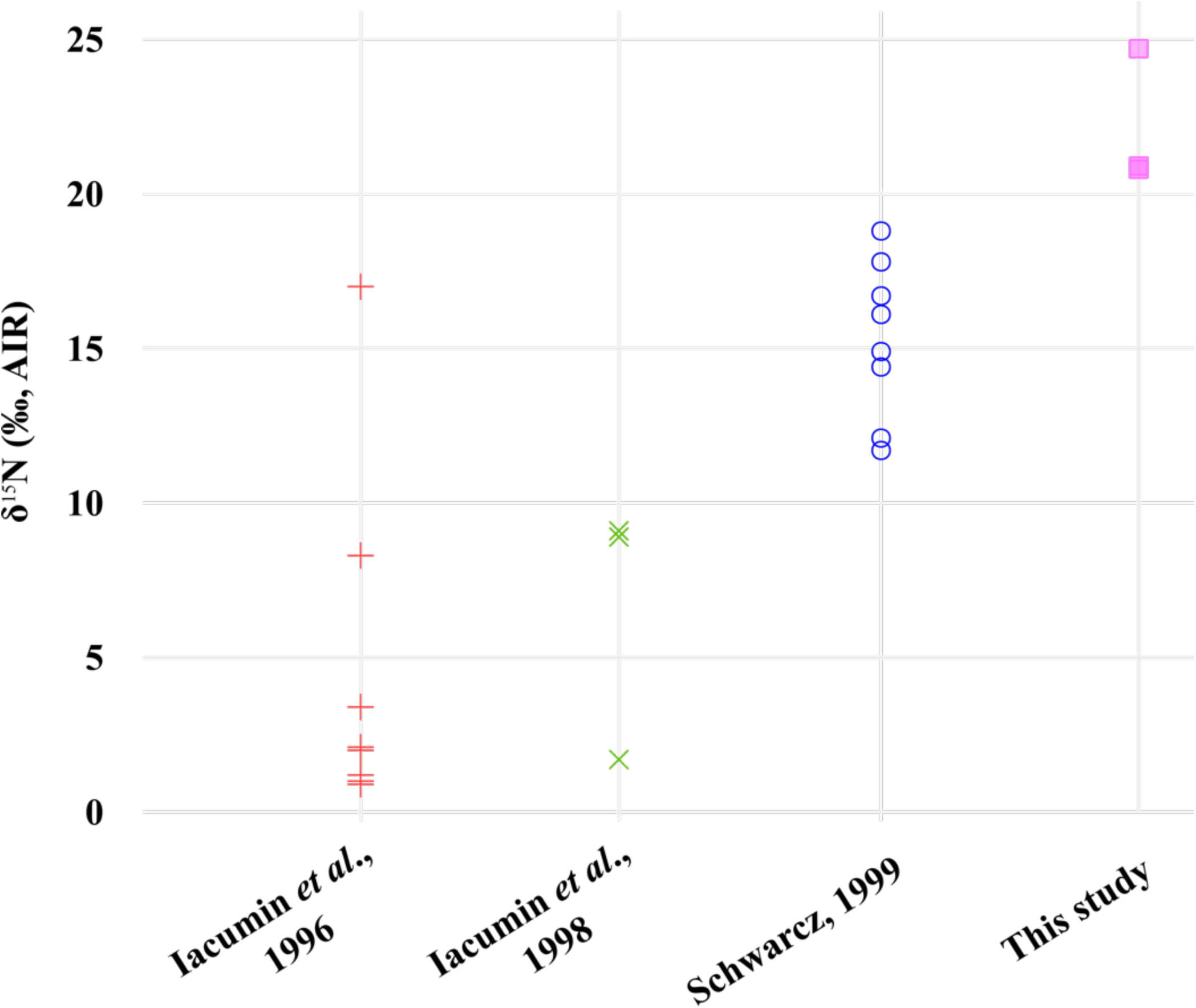
Nitrogen isotope compositions (δ^15^N) of the plant remains of our study compared with previous published data for modern and archaeobotanical remains (Iacumin et al., 1998, 1996; Schwarcz et al., 1999).

The Predynastic mummies in our study were discovered in the Nile valley, in an environment that was less subject to aridity, especially for the Predynastic Period, that was more humid than the Dynastic period (Touzeau et al., 2013). Moreover, the active floodplain in the vicinity of Thebes, ancient Luqsor, near our archaeological study sites, was active for a considerable distance around the Nile in Predynastic times (Peeters et al., 2024). Even though the lack of annual flooding of the Nile may have contributed to an increase in local soil aridity (Hassan and Graham-Campbell, 1997; Welc and Marks, 2014), this phenomenon alone cannot explain the δ^15^N values observed.

### 6. Alteration impacting on δ^15^N values?

We can speculate that the observed ^15^N enrichment could be attributed to a post-burial phenomenon, such as microbial or thermal alteration, given that these kinds of alterations might change δ^15^N values of tissues. This phenomenon would have taken place after burial but before the desiccation of the tissues, and all matrices would be expected to undergo similar changes. Certain types of alteration, in particular bacterial alteration, have shown that collagen can be enriched in ^15^N compared to its original composition (Balzer et al., 1997), whereas thermal alteration could slightly reduce δ^15^N values (Harbeck and Grupe, 2009). However, the δ^15^N values of the skins do not exhibit a significant divergence from the δ^15^N values reported for other skins in the literature (Iacumin et al., 1998, 1996; Johns, 2012) and while an enrichment is notable, it is less pronounced than that observed in plants. We therefore rule out this hypothesis.

### 7. Soil fertilisation hypothesis

The soil of the Nile Valley was naturally fertilised by the annual floods of the Nile, which deposited nutrient-rich silt and alluvium (e.g., phosphorus) onto the floodplains, resulting in a natural fertilisation of the soil (El-Ramady et al., 2013; Hughes, 1992). However, fertilisation with Nile silts alone is not sufficient to explain the high δ^15^N values, as previous studies from this period did not report such elevated values in plants or animals (e.g. Iacumin et al., 1998, 1996; Touzeau et al., 2014). We suggest that the elevated δ^15^N values observed in this study may be attributable to soil fertilisation practices. The practice of manure fertilisation has been documented in Europe since the Neolithic (Bogaard et al., 2013), and δ^15^N values in soils and plants vary depending on the type of fertiliser. For instance, the application of guano or pig manure can raise plant δ^15^N levels by up to +19‰ (Szpak, 2014; Szpak et al., 2014, 2012), while cattle manure tends to have a more moderate effect (Szpak, 2014). Another method, known as middening, where household waste is dumped onto the soil, has also been used as a soil amendment, as documented at Tell Sabi Abyad in northern Mesopotamia, dating back to the 7th millennium BC (Styring et al., 2017). If soil fertilisation occurred in the study area, it implies that such practices were long-standing, as it takes several years for isotopic shifts in plant δ^15^N to manifest following manure application (Fraser et al., 2011). Although manure fertilisation has been hypothesised for Ancient Egypt (Wenke, 2009), no direct evidence has been recorded to date. Our findings strongly suggest that soil fertilisation, possibly with manure, was already in use during the Predynastic Period in this region.

While ^15^N-enrichment is expected to propagate through the food chain, the plants associated with the mummified remains are non-edible. It would be unlikely for these populations to selectively fertilise specific non-edible plants and not others. It is more plausible that all local plants, including those consumed, were similarly fertilised, which would lead to elevated δ^15^N values throughout the trophic chain. However, here, plants are significantly more enriched in δ^15^N than animal or human tissues. This suggests that humans and animals may not have lived right next to the locations where the associated plants grew.

## Conclusion

Multi isotopic analysis of archaeological remains from Predynastic Egypt provides us a more precise understanding of the local farming methods employed during this period. Three of the four mummies in this study were buried enveloped in textiles, a plant mat and animal skins and one of them was only enveloped in plant material. Although the degree of preservation of human samples analysed in this study is sub-optimal, the joint analyses of plants, animal skins, and human skins have revealed high δ^15^N values observed across all tissues studied. Moreover, the results are contextualised within the broader framework of available data from the same period, as well as data from Dynastic Period, providing a better understanding of the findings and showing that our results are in line with the literature. High δ^15^N values measured in our study cannot be alone explained by aridity, water stress or consumption of Nile fish as previously proposed in former studies (e.g., Touzeau et al., 2014). Therefore, we suggest the possibility of local fertilisation practices involving the use of manure from livestock in Predynastic Egypt, a hypothesis that has likely been underestimated to date and that merits further investigation as part of a more comprehensive study. If such a hypothesis was to be confirmed in the future by direct archaeological evidence, it would imply its use in a somewhat extended period, because an occasional soil amendment would not enrich the tissues in ^15^N studied to such an extent. Although existing literature provides values similar to ours, the analyses conducted on plants and animal skins in this study have led to different conclusions from those previously reported.

This study highlights the importance of analysing environmental elements associated with mummies, demonstrating how these materials can provide valuable insights into past ecological conditions and human-environment interactions. Implementing this comprehensive approach more systematically across museum collections could yield significant discoveries and enhance our understanding of ancient societies. This study also shows the need for a more comprehensive examination of a tissue, the skin, which is currently understudied due to its scarcity at archaeological sites. A more systematic investigation of various human tissues could prove beneficial.

## Acknowledgments

These mummies were studied in a larger program called Human Egyptian LYon COnfluences Mummies (HELYCOM) – Mourir pour renaître. We would like to thank the staff of the Musée Confluences in Lyon, France, who allowed us to sample their material on numerous occasions. In particular, we would like to thank François Vigouroux for his assistance and guidance during the sampling process. We would like to thank the ENS Lyon for letting us use their freeze-dryer, and especially Salomé Ansanay-Alex for her help. We would also like to extend our gratitude to Victoria Asensi-Amoros of Xylodata, Paris, for her assistance in plant identification, and to Laurianne Robinet, Sylvie Heu-Thao, Sophie Cersoy of the Centre de Recherche sur la Conservation, MNHN, Paris, for their contributions to the identification of animal skins, and to Michel Billard, paleopathologist. Finally, we also thank Eric Crubézy, Université Toulouse III-Paul Sabatier for his help and Yvan Tristant, Katholieke Universiteit Leuven, for his comments about scientific data related to agricultural practices in Predynastic Egypt

## Author Contributions

EP, AP, JPF and CL conceived and designed the study. Material was sampled by EP, DB, and CL on the expert advice of AP. Preparation of the samples was made by EP and AVL. Isotope analyses were performed by EP, AVL, and FF. Radiocarbon dating was performed by PR. The first draft of the manuscript has been written by EP, JPF and CL, and all authors commented on previous versions of the manuscript. All authors read and approved the final manuscript.

## Data availability

All data are available in the main text or in the Supplementary Material file.

